# The soursop genome and comparative genomics of basal angiosperms provide new insights on evolutionary incongruence

**DOI:** 10.1101/639153

**Authors:** Joeri S. Strijk, Damien D. Hinsinger, Mareike M. Roeder, Lars W. Chatrou, Thomas L. P. Couvreur, Roy H. J. Erkens, Hervé Sauquet, Michael D. Pirie, Daniel C. Thomas, Kunfang Cao

## Abstract

Deep relationships and the sequence of divergence among major lineages of angiosperms (magnoliids, monocots and eudicots) remain ambiguous and differ depending on analytical approaches and datasets used. Complete genomes potentially provide opportunities to resolve these uncertainties, but two recently published magnoliid genomes instead deliver further conflicting signals. To disentangle key angiosperm relationships, we report a high-quality draft genome for the soursop (*Annona muricata*, Annonaceae). We reconstructed phylogenomic trees and show that the soursop represents a genomic mosaic supporting different histories, with scaffolds almost exclusively supporting single topologies. However, coalescent methods and a majority of genes support magnoliids as sister to monocots and eudicots, where previous whole genome-based studies remained inconclusive. This result is clear and consistent with recent studies using plastomes. The soursop genome highlights the need for more early diverging angiosperm genomes and critical assessment of the suitability of such genomes for inferring evolutionary history.

## Introduction

Reconstructing the sequence of rapid speciation events in deep time is a major challenge in evolutionary inference. Bursts of diversification result in short branches within phylogenetic trees and the persistence of discrepancies between histories of individual genes, genomes and the underlying species tree (Oliver et al. 2013). These phenomena are prevalent across the Tree of Life, including the origin of important, species-rich lineages such as tetrapods (Song et al. 2012; Jarvis et al. 2014), insects (Freitas et al. 2018), and flowering plants (Tank et al. 2015). Whole genome sequencing and application of the multispecies coalescent offer a new, unprecedented opportunity to reconstruct such recalcitrant relationships (Edwards 2009; Edwards et al. 2016).

The emergence of flowering plants (angiosperms) was a geologically sudden and ecologically transformative event in the history of life. Recent analyses have suggested that the major angiosperm clades diverged in quick succession within the Cretaceous, with monocots, magnoliids and eudicots starting to diversify within ca 5 Ma (Sauquet and Magallón 2018).

Despite steady progress in the reconstruction of angiosperm phylogeny and evolution^e.g.6^, deeper nodes have proven notoriously difficult to resolve, in particular the relationships between *Ceratophyllum*, Chloranthaceae, monocots, magnoliids, and eudicots (Ruhfel et al. 2014; Wickett et al. 2014; Zeng et al. 2014).

Potential relationships among the main angiosperms lineages can be summarized as either 1) magnoliids sister to eudicots + monocots (Moore et al. 2010; Qiu et al. 2010; Soltis et al. 2011; Zhang et al. 2012; Magallón et al. 2015); 2) magnoliids sister to eudicots (Bell et al. 2010; Moore et al. 2011; Zeng et al. 2014); or 3) magnoliids sister to monocots (Nickrent and Soltis 2006). Each of these topologies were inferred from organelles (chloroplast and mitochondrial loci) and/or nuclear datasets, with variable levels of support depending the analytical method and taxonomic sampling used. In parallel, recently published studies challenge our understanding of the early evolution of angiosperms. Reconstruction of the ancestral angiosperm genome shows a reduction in the number of chromosomes between the divergence of the putative sister taxa of all remaining angiosperms *Amborella* and the most recent common ancestor (MRCA) of eudicots (Murat et al. 2017). In addition to this assessment of early angiosperm genomic features, Sauquet *et al*. (Sauquet et al. 2017) suggested that the early evolution of angiosperm flowers was marked by successive reductions in the number of whorls of both the perianth and the androecium. In both cases, disparate reductions may have paved the way for the evolution of clade specific features of genomes and flower morphology in contemporary clades.

Since the publication of the first plant genome, that of *Arabidopsis thaliana* (Initiative 2000), there has been a steady increase in the number of sequenced eudicot and monocot genomes. However, with the exception of the iconic *Amborella trichopoda*, basal angiosperm diversity represented by the ancient lineages of Nymphaeales, Austrobaileyales, Chloranthales, and magnoliids has largely been overlooked. After eudicots and monocots, Magnoliidae are the most diverse clade of angiosperms (Massoni et al. 2014) with 9,000-10,000 species in four orders (Canellales, Piperales, Laurales and Magnoliales). However, despite this diversity and economic value (e.g. avocado, black pepper, cinnamon, soursop), only two genomes have been published to date (Chaw et al. 2019; Chen et al. 2019). Analysis of such genomic data was expected to resolve the still unclear relationships of magnoliids with the rest of angiosperms (Soltis and Soltis 2019). However, the results strongly disagreed on the position of magnoliids, supporting either a sister relationship to eudicots and monocots (Chen et al. 2019), or to eudicots alone (Chaw et al. 2019). This disagreement could be an analytical artefact caused by whole genome duplication (WGD) and chromosomal rearrangements as apparent in both *Liriodendron chinense* (the Chinese tulip poplar) and *Cinnamomum kanehirae* (the stout camphor tree) after their divergence from monocots or eudicots. More magnoliid genomes and critical assessment of the potential for such phenomena to impact phylogenetic inference is needed to break the impasse.

We sequenced the genome of *Annona muricata* (the soursop) which is one of the c. 2450 species of the custard apple family (Annonaceae) (Rainer and Chatrou 2014), the second most species rich family of magnoliids (Chatrou et al. 2012). Its species are frequent components of tropical rain forests worldwide (Gentry 1993; Tchouto et al. 2006; Punyasena et al. 2008; Sonké and Couvreur 2014). Widely known examples include ylang-ylang (*Cananga odorata*), used for its essential oils, and species of the Neotropical genus *Annona*, such as the soursop, cultivated for their edible fruits.We then undertook comparative intergenomic analyses in magnoliids, reconstructed the relationships among the three major lineages of angiosperms and explored the gene tree incongruence patterns during early angiosperm diversification.

## Results

To assemble the genome of the soursop, we extracted DNA from a soursop tree (*Annona muricata*) grown in Xishuangbanna Tropical Botanical Garden (XTBG, Menglun, China), and generated 131.43 (164.47x) and 36.95 Gb (46.24x) of Illumina and PacBio reads, respectively (Supplementary Table 1). SOAPdenovo 2.04 was used to assemble the first draft of the genome, reaching a contig N50 of approximately 700 kb.

The soursop assembly was further refined using both 10X Genomics data (180.04 Gb – 225.30x) and Bionano optical mapping data (95.9 Gb – 120.01x) (Supplementary Table 1) to improve the scaffold N50 to 3.4 Mb, the longest scaffold being 20.46 Mb (GC content of 34.35%, Supplementary Table 2). The total assembly length of the soursop genome was 656.78Mb (scaffold assembly). Scaffolds longer than 100kb totaled 646.64Mb (98.45% of the total length). This level of contiguity is similar to the one obtained in *Liriodendron chinensis* (N50=3.5 Mb (Chen et al. 2019)) but smaller than obtained in *Cinnamomum kanehirae* (N50=50.4Mb after Hi-C scaffolding (Chaw et al. 2019), and better than other genomes assembled at scaffold-level (e.g. (Yang et al. 2018; Arimoto et al. 2019; Zhang et al. 2019)). A total of 444.32 Gb of data were produced using Illumina, PacBio, 10X Genomics and Bionano technologies, corresponding to 556x coverage of the soursop genome. We assessed the quality of the soursop genome assembly by mapping back the Illumina reads against the assembly. A total of 97.16% reads can be mapped, covering >99.92% of the genome, excluding gaps. 99.81% of the genome was covered with a depth >20x, which guaranteed the high accuracy of the assembly for SNPs detection (Supplementary Table 3). The final assembly comprised 949 scaffolds, 29 greater than 5Mb in length. SNP calling on the final assembly yielded a heterozygosity rate of 0.032%, lower than 0.08% as estimated by the K-mer analysis (Supplementary Fig. 1).

Repeats accounted for 54.87% of the genome (Supplementary Table 4, Fig. 1b) and were masked for genome annotation. It is slightly less than in *Cinnamomum* (Lauraceae, 48%) and *Liriodendron* (Magnoliaceae, 61.64%). Long Terminal Repeat (LTR) retrotransposons were the most abundant, representing 41.28% of the genome (56.25% in *L. chinense*), followed by DNA repeats (7.29%). The stout camphor tree genome exhibited a different balance between types, with LTR (25.53%) and DNA transposable elements (12.67%) being less dominant. No significant recent accumulation of LTRs and LINEs was found in the interspersed repeat landscape, but a concordant accumulation around 40 units was detected (Fig. 1c). Assuming a substitution rate similar to the one found in *Liriodendron* (1.51×10-9 subst./site/year), we estimate this burst of transposable elements to have occurred 130-150 Ma ago. By far the main contributor to this old expansion of repeat copy-numbers were the LTRs, with an increase of up-to approximately 1% at 42 units.

**Figure 1.**
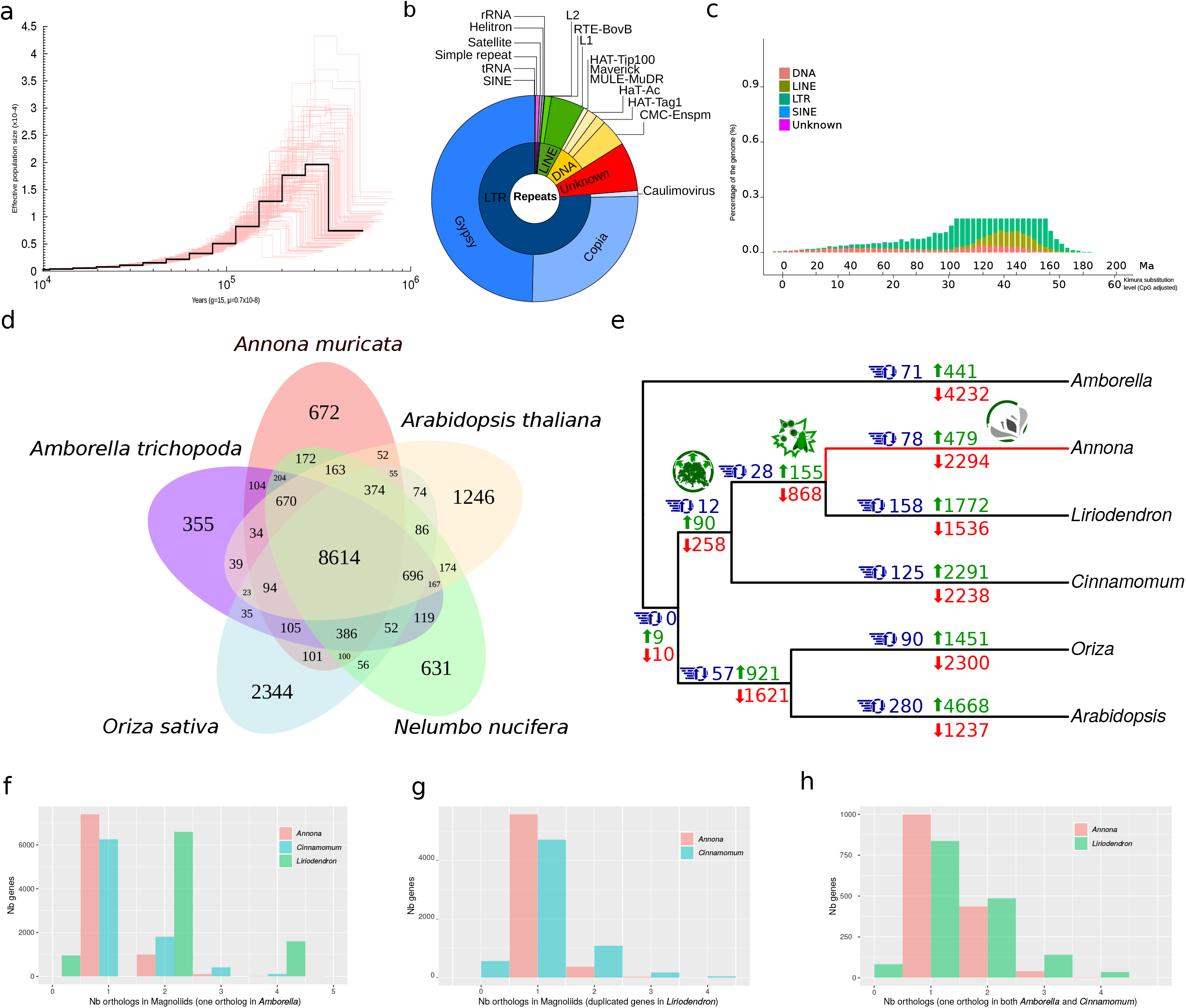
Characteristics of the soursop genome and comparative analyses.(a) Effective population size history inferred by the PSMC method (black line), with one hundred bootstraps shown (red lines. (b) Distribution of repeat classes in the soursop genome. (c) Divergence distribution of transposable elements in the genome of *Annona muricata*. Both Kimura substitution level (CpG adjusted) and absolute time are given. (d) Venn diagram of shared orthologous gene families in *Amborella trichopoda, Arabidopsis thaliana, Nelumbo nucifera, Oriza sativa* and *Annona muricata*, based on the presence of a representative gene in at least one of the grouped species. Numbers of clusters are provided in the intersections. (e) Coalescent tree of the dataset comprising the three magnoliid genomes plus *Amborella* and representative of eudicots (*Arabidopsis*) and monocots (*Oriza*), based on 1578 orthologs and ASTRAL-III reconstruction. Number of CAFE-reconstructed gene families variation are shown on the branches (green: expansion; red: contractions; blue: rapid changes). Major annotations experiencing rapid expansion on magnoliids branches are pictured (see main text for details). (f) Number of families (vertical axis) according to the number of orthologs (horizontal axis) found in the genome of *Annona* (red), *Liriodendron* (green) and *Cinnamomum* (blue) for families containing one single ortholog in *Amborella*. (g) Number of families (vertical axis) according to the number of orthologs (horizontal axis) found in the genome of *Annona* (red) and *Cinnamomum* (blue) for families containing two orthologs in *Liriodendron*. (h) Number of families (vertical axis) according to the number of orthologs (horizontal axis) found in the genome of *Annona* (red) and *Liriodendron* (green) for families containing one single ortholog in both *Amborella* and *Cinnamomum*.

Genes were annotated using a comprehensive strategy combining *ab initio* prediction with protein homology detection and transcriptomic data from leaves, flowers, bark and both young and ripening fruits (Supplementary Table 1). We combined the gene models predicted by these three approaches in EvidenceModeler v1.1.1 (EVM) to remove the redundant gene structures and filtered the resulting gene models by removing (1) the coding regions ≤150bp and (2) models supported only by *ab initio* methods and with FPKM<1. We identified 23,375 genes and 21,036 genes supported by at least two methods (Supplementary Fig. 3), with an average coding-region length of 1.1 kb and 4.79 exons per gene, similar to other angiosperms (Supplementary Table 5). 22,769 (97.4%) of these genes were annotated through SwissProt and TrEMBL and GO-terms were retrieved for 20,595 (88.1%) genes (Supplementary Table 6). We assessed both the quality of our gene predictions and completeness of our assembly using BUSCO and CEGMA approaches. 231 CEGs genes (93.15%) and 899 (94%) of the BUSCO orthologous single copy genes were retrieved from the soursop assembly (Supplementary Table 7).

We determined *A. muricata* exhibits heterozygous and homozygous SNP ratios of 0.0032% and 0.0001%, respectively. This very low heterozygosity, usually found in cultivated species that experienced strong bottlenecks during domestication (Eyre-Walker et al. 1998; Doebley et al. 2006; Zhu et al. 2007), was not due to an intense, recent decrease in population size, as shown by our PSMC analysis. Instead, the very low heterozygosity observed in soursop was rather due to a slow and regular reduction of the species population sizes (Fig. 1a).

By assembling transcriptomes from several organs, in addition to homology-based prediction and *de novo* predictions, we identified 21,036 genes supported by at least two methods (Supplementary Fig. 3). We assessed both the quality of our gene predictions and completeness of our assembly using BUSCO and CEGMA approaches. 231 CEGs genes (93.15%) and 899 (94%) of the BUSCO orthologous single copy genes were retrieved from the soursop assembly (Supplementary Table 7).

To infer phylogenetic relationships among major clades of angiosperms, we compared the soursop genome with the genomes of eleven other species. We included *Amborella trichopoda*, the putative sister lineage to all extant angiosperms, selected representatives from monocots (*Oryza sativa, Musa acuminata*), and key lineages of eudicots, including Ranunculales (*Aquilegia coerulea*), Proteales (*Nelumbo nucifera*), superrosids (*Vitis vinifera, Quercus robur, Arabidopsis thaliana*), and superasterids (*Amaranthus hypochondriacus, Helianthus annuus, Coffea canephora*). We used *all-against-all* protein sequence similarity searches with OrthoMCL to identify 398,668 orthologs with at least one representative in angiosperms. 672 of these were unique to *A. muricata* and 8,614 were found to be shared with four species representative of other main lineages of angiosperms (see Fig. 1d).

To investigate patterns of incongruence and support in the reconstructed phylogeny of the main angiosperms lineages, we focused on testing three main hypotheses and corresponding topologies: 1) magnoliids sister to eudicots + monocots [(*Annona*, (eudicots, monocots), hereafter referred to as SAT]; 2) magnoliids sister to eudicots [(monocots, (*Annona*, eudicots)), SMT)]; and 3) magnoliids sister to monocots [(eudicots, (*Annona*, monocots), SET].

We assessed conflicting signals in the genome of the soursop (Fig. 2), using three dataset. Firstly, we used a set of 2426 orthologs identified in the quartet (*Annona muricata, Arabidopsis thaliana, Amborella trichopoda* and *Oryza sativa*), maximising the number of loci. Secondly, a set of 689 orthologs identified in the 12 species was used for comparative analyses as described above, maximizing the number of species. Finally, a set of 1578 orthologs identified in the previous quartet, plus *Cinnamomum kanehirae* and *Liriodendron chinense*, was used maximizing the magnoliid representation.

**Figure 2.**
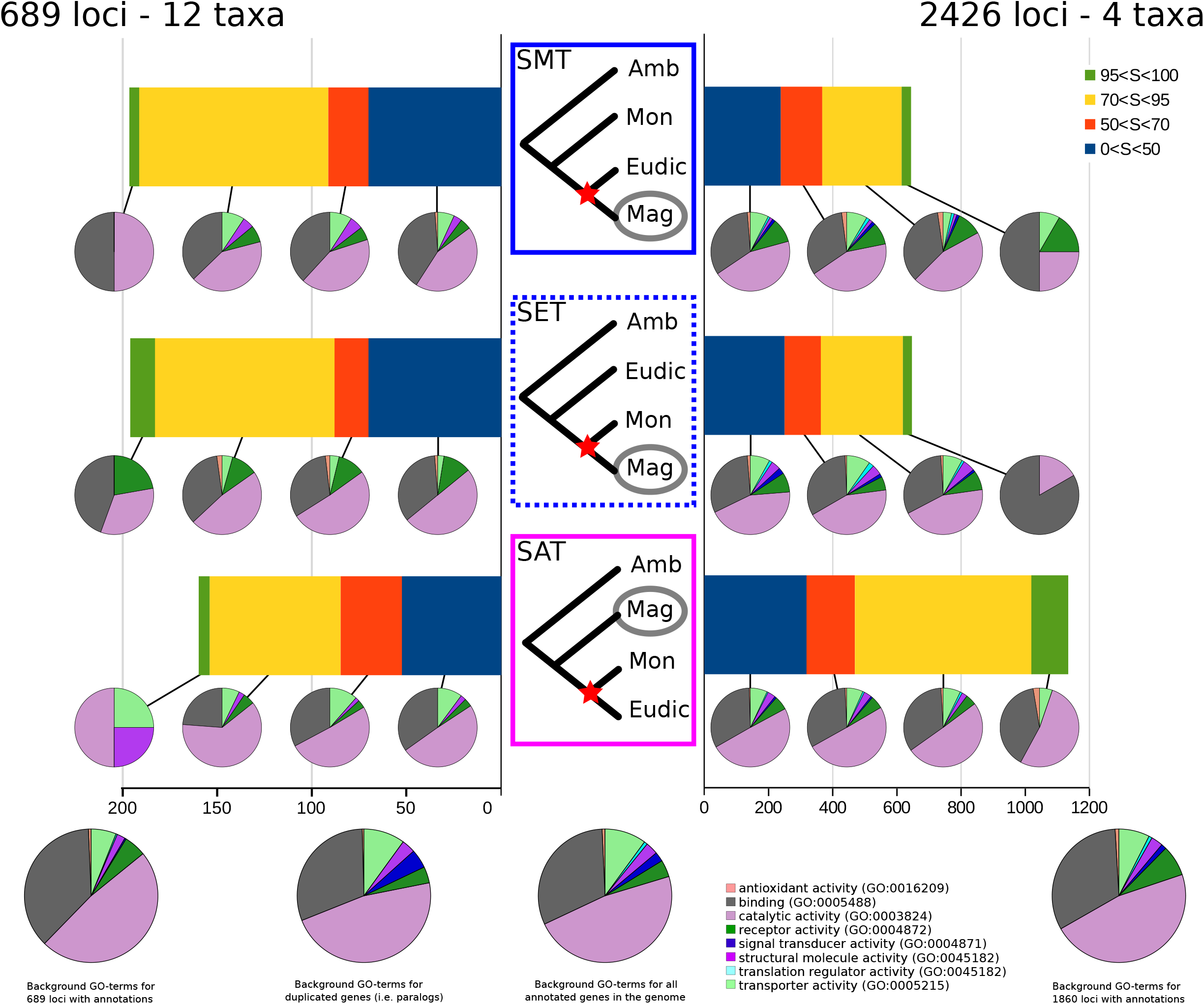
Incongruence in the phylogenetic signal in the genome of *Annona muricata*. Left: using 689 orthologous loci found in 12 angiosperm species. Right: using 2426 orthologous loci found in 4 species representative of major clades in angiosperms. For each of the topologies shown in the center, the genes supporting this topology are sorted by the support of the nodes indicated by stars. The GO-terms associated with each category of support for each topology are indicated as pie-chart below the histogram. Background GO-terms distribution for 689 loci (left), 2426 loci (right) and the total annotated genes in *A. muricata* (center) are shown below the graph. Topology supported by the concatenated 689 loci, coalescent analysis of 689 loci and by the both concatenated and coalescent 2426 loci are highlighted in solid blue, dashed blue and solid pink, respectively.

Using the set of 2426 orthologs and single-gene ML phylogenetic reconstructions, we show that the SMT topology found using *Cinnamomum* as a representative of magnoliids, is only supported by ~16.8 % of the genes (29) with a clear evolutionary signal (SH-like values >0.95 for one topology) (SET: 16.27% - 28 genes; SAT: 66.9% - 115 genes). When taking into account the slightly weaker phylogenetic signal (SH-like values >0.70), the majority (54.3 %) of the 1228 genes supported SAT (SET: 23%; SMT: 22.5%) (Supplementary Table 8). Assuming gene tree differences are the result of coalescent stochasticity, we performed coalescence-based analyses (STAR, NJst, MP-EST, ASTRAL-III) of the three datasets to reconstruct a species tree (Supplementary Fig. 4). Using 689 loci (12 taxa), all three coalescent methods (STAR, NJst, MP-EST) retrieved a SET topology, whereas both the 2426 loci (4 representative taxa) and the 1578 loci dataset (4 representative taxa + 2 published magnoliids) dataset retrieved a SAT topology. Interestingly, in both the 689 and 2426 loci datasets, the branch supporting the position of *Annona* in the trio *Annona-*-monocots-eudicots is very short (Supplementary Fig. 4), suggesting that divergence of the major clades occurred within a short time frame.

The rise of angiosperms during the Cretaceous was likely triggered by their interaction with pollinators and the early onset of morphological adaptations in the group (e.g. reproductive and vegetative parts)^20^, suggesting that the underlying molecular functions (such as those linked to flowering processes or adaptation to insect herbivory) could have quickly diversified in response, and could thus reflect diversification patterns and ecological adaptation, instead of phylogeny. To identify potential functions triggering bias in the evolutionary signal contained in the soursop genome, we compared the GO-terms annotations for molecular functions in the genes supporting the different topologies (SH-like support >0.95) highlighting different trends in functional annotations according to the topology they supported (P-value < 0.001, Friedman rank sum test, see Supplementary Fig. 5). Briefly, differences among topologies were significant only for Catalytic Activity (p<0.005, Kruskal-Wallis rank sum test) and Receptor Activity (p<0.05). We found the “Receptor activity” genes were over-represented (p<0.005) in the SMT topology (when considering genes giving topologies without support (0<SH<50) relative to the background.

To investigate the genomic landscape of incongruent phylogenetic signals, we compared scaffold features in terms of ortholog number and density (Supplementary Fig. 6) for the 2426 loci dataset. The 181 scaffold containing orthologs (19%) included an average of ~22 orthologous coding regions (max=799, found in the scaffold_1, see Supplementary Table 10), with a density of 1.33 x10^−2^ orthologs per kb. No correlation was found between the length of the scaffold and the proportion of genes supporting a given topology (Supplementary Fig. 6), excluding a potential bias toward a given topology during assembly. However, the median of the proportion of genes in a scaffold supporting the SAT was close to 23%, instead of 4% for both the SMT and SET (Supplementary Fig. 7, box plots). Considering a given topology, density was one or two orders of magnitude lower, the genes supporting the SET, the SMT and SAT hypotheses being found with densities of 5.09 x10^−4^, 2.13 x10^−3^ and 6.34 x10^−4^ genes per kb, respectively. Compared to the density of all orthologs found in each scaffold (aka “the background”), the highest relative density was found in the genes supporting the SAT (¼ of the background density), followed by the SMT and SET hypotheses, with 13.9% and 10.6% respectively. We generated heatmaps of the occurrences of orthologs and showed that distribution of orthologs is uneven across scaffolds: scaffolds rich in orthologs supporting a given topology did not contain a significant number of orthologs supporting a conflicting topology (Supplementary Fig. 6). Most of the scaffolds contained few orthologs supporting a given topology (Supplementary Fig. 6a), with only a few scaffolds showing a high topology-supporting ortholog density (Supplementary Fig. 6b). Altogether, our results strongly support the magnoliids as sister to a clade containing (eudicots+monocots), i.e. the SAT topology (Fig. 3).

**Figure 3.**
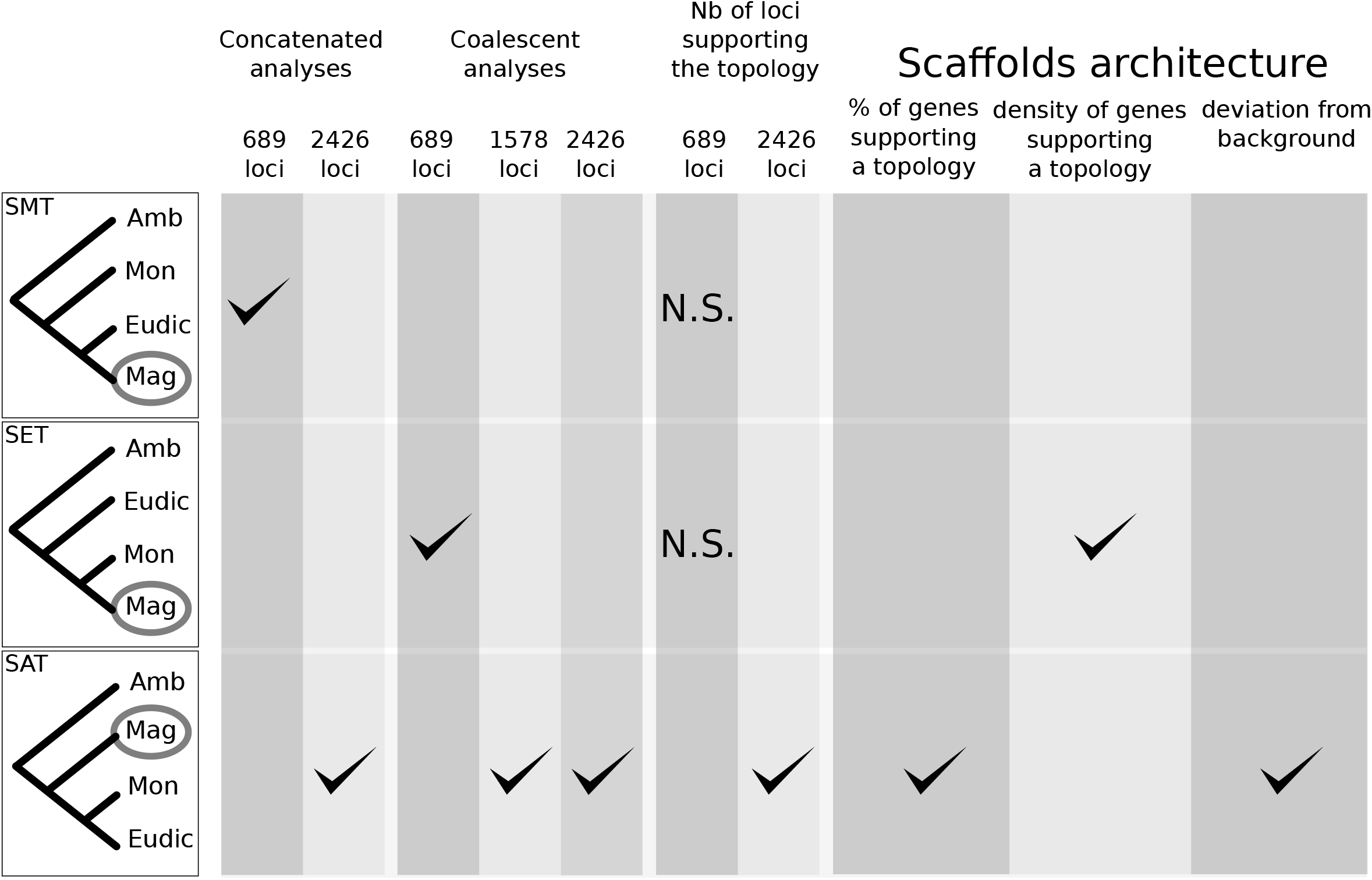
Summary of the supportive evidence for each tested topology. N.S.: non-significant results.

Comparing gene content in *Annona* with that found in the stout camphor tree, we found a striking difference in diversity of resistance genes. Of 387 resistance genes in *Cinnamomum*, 82% were nucleotide-binding site leucine-rich repeat (NBS-LRR) or with a putative coiled-coil domain (CC-NBS-LRR). By contrast, the soursop genome contains a similar number of resistance genes (301 annotations), but only 0.66% (2 genes) of them are NBS-LRR or CC-NBS-LRR genes. These results suggest the presence of different evolutionary strategies within magnoliids with respect to pathogen resistance. We explore the expansion of gene families in magnoliid lineages by adding *Cinnamomum* and *Liriodendron* to the quartet (*Annona muricata, Arabidopsis thaliana, Amborella trichopoda* and *Oryza sativa*) (Fig. 1e). GO-terms from annotations of these gene families show that the lineage of *Annona* experienced a fast expansion of both the MAD1 protein family (+6 copies, involved in flowering time -GO:0009908- and cold adaptation -GO:0009409), and metabolism through mitochondrial fission (GO:0000266), regulation of transcription (GO:0006383, GO:0006366, GO:0045892) and organism development (GO:0007275). Half of the expanded gene families with annotations in the branch of Magnoliales (*Liriodendron, Annona*) were involved in disease or pathogens resistance. On the contrary, gene families experiencing fast expansion on the magnoliids branch ((*Liriodendron, Annona),Cinnamomum*) were mainly involved in growth function, like cell wall biogenesis (GO:0042546), membrane fission (GO:0090148) and metabolism of peptides (GO:0006518), proteins (GO:0046777) or mitosis cytokinesis (GO:0000281). Notably, gene family expansion is consistently lower along internal branches (approximately 1/10^th^ of the gene family expansion is found on their sister branches). Whole genome duplications (WGD) are suspected to be a significant trigger factor in the diversification of angiosperms (Tank et al, 2015; Vamosi et al, 2018). Ks distribution of both the soursop paralogs (Supplementary Fig. 2a) and synteny analysis using i-adhore 3.0 and SynMap as implemented in CoGe (Lyons & Freeling, 2008; Lyons et al., 2008) did not reveal any obvious pattern of recent tandem- or whole-genome duplication. However, an old duplication (around 1.5 Ks units) was found in the soursop genome. This contrasts with recent studies in magnoliids (Chaw et al. 2019; Chen et al. 2019), where the authors found WGD with lower Ks values (thus potentially more recent), and hypothesized them to be shared in magnoliids and thus older than the divergence between Lauraceae and Magnoliaceae.

In order to assess whether the events identified in *Cinnamomum* and *Liriodendron* correspond to a magnoliids-shared WGD, or to independent events, we compared the syntenic graph of *Arabidopsis* and each of the magnoliids genomes. In each magnoliid genome, we found evidence of duplicated syntenic blocks (stronger in *Liriodendron*, but inconclusive in *Cinnamomum* depending on whether one or two WGD occured).

We evaluated the distribution of gene copy numbers among magnoliids for gene families with only one copy in *Amborella* (i.e. the number of copies found in each magnoliid for gene families in which *Amborella* has only one copy). We found that despite a WGD event reported in *Liriodendron* and two in *Cinnamomum*, only the former displayed two gene copies for the majority of the gene families, with *Annona* holding one gene copy for almost all families (Fig. 1f). Conversely, *Annona* and *Cinnamomum* contained only one copy of almost all the gene families for which *Liriodendron* contains two copies (i.e. the gene families that show a WGD signal, Fig. 1g). For gene families in which only one copy was found in both *Amborella* and *Cinnamomum*, both *Annona* and *Liriodendron* showed a similar pattern of mainly unique orthologs families, with about half of the families being duplicated in both species (Fig. 1h). However, the ratio duplicated/unique occurrence in orthogroups was smaller in *Annona* (0.43) than in *Liriodendron* (0.58), suggesting a shared ancestral WGD in magnoliids. We used MCscan to detect syntenic regions in magnoliids and *Amborella*, and detected large one-copy portions of the genome in *Amborella* occurring as duplicates in *Liriodendron*, as expected from the previous studies (Chen et al. 2019) and our results above. More surprisingly, duplicated syntenic regions in *Cinnamomum* showed evidence for two rounds of WGD (i.e. four copies of a single *Amborella* syntenic region), in contradiction with our results above, but according to previous studies. However, after careful evaluation of the synteny graphs, we found limited evidence (i.e. few/shorter duplicated syntenic regions) of WGD in the genome of the soursop compared to *Amborella*. To characterize more precisely the ages and distributions of these duplication events, we analyzed the Ks distribution curves for the paranomes (i.e. the complete set of paralogs in a genome) of the soursop and the two published magnoliids. We used WGD because it has been shown that node-averaged histograms are more accurate than weighted ones to infer ancient WGD (Tiley et al. 2018).

Contrary to other magnoliids, the paranome of the soursop (Supplementary Fig. 2a) did not show the usual acute peak corresponding to newly duplicated genes that are continuously generated by small scale duplications events (e.g. tandem duplications), but showed an older small scale duplication event peak. No realistic Gaussian mixture model (i.e. implying <4 WGD) was selected by either the BIC or AIC criterion, but the Δ_BIC_ and Δ_AIC_ favored 2 component models for *Annona* and *Cinnamomum*, with a less clear signal in *Liriodendron* (Supplementary Fig. 2a,b,c). In addition to the standard GMM method, we also used the BGMM method and confirmed that fitting more complex Gaussian models only resulted in components of negligible weights (results not shown). Considering two components for *Annona* and *Liriodendron*, and two-three components for *Cinnamomum* (as indicated by the results above), the main peak in *Annona* was found around 1.3-1.5 Ks units, whereas it occured around 0.8 Ks units (two components) or 0.4 and 1.3-1.4 Ks units (three components) in *Cinnamomum* (partially comparable - for the two components - with results from previous studies (Chaw et al. 2019), as we did not identify a peak at 0.76). Notably, the oldest peak in *Cinnamomum* located approximately at the same divergence as the peak in *Annona*, suggesting a potentially shared WGD. We identified a peak at 0.6 Ks units in *Liriodendron*, compatible with the location of the youngest peak in *Cinnamomum*, but younger than previously inferred (Chen et al. 2019). Considering the divergence of paralogs in each of the magnoliids, it seems unlikely that they share a common WGD event. Indeed, the pattern of potentially shared WGD (*Annona+Cinnamomum* ~1.5 Ks units; *Liriodendron* + *Cinnamomum* ~0.5 Ks units) seems incompatible with both the current and our reconstructed hypothesis of magnoliids evolution.

By performing one-vs-one ortholog comparisons in magnoliids, we found that the divergence of the *Annona* and *Liriodendron* lineage occured around 0.6-0.7 Ks units (Supplementary Fig. 2d), while the divergence between Magnoliales and Laurales appeared to be slightly older at 1.0-1.1 Ks units (Supplementary Fig. 2e). The divergence of magnoliids (represented by *Annona*) from *Amborella* took place at 1.8-1.9 Ks units (Supplementary Fig. 2f). This confirms the likely absence of a shared WGD in magnoliids, and places the WGD events observed in both *Cinnamomum* and *Liriodendron* subsequent to their MRCA with *Annona*.

## Discussion

Increasing availability of high quality genome assemblies enables far greater insight into challenging phylogenetic problems, like ancient and rapid diversification events, as epitomised by the early evolutionary history of angiosperms. The soursop genome provides evidence for the rapid sequential divergence of magnoliids, monocots and eudicots, with a mosaic of phylogenetic signals across the genome reflecting coalescence and potentially hybridisation between closely related ancestral lineages. This was followed by relative structural stasis since the Jurassic-Cretaceous boundary, with none or very few ongoing small scale duplications, fewer paralogs than other magnoliids, no significant burst of transposable elements and few expanded gene families along the branches leading to *Annona*. To our knowledge, the soursop is the first nuclear genome displaying such extensive signs of “fossilization” [notwithstanding the mitochondrial genome of *Liriodendron tulipifera* which has been also described as “fossilized” (Richardson et al. 2013)]. A genome which has retained the original characteristics of the ancestral magnoliid lineage, will be an invaluable resource for future studies on angiosperm early diversification. The apparent stability of the *Annona* genome over time is notable given that gene family expansion (such as in (Chaw et al. 2019)),increase of transposable elements (Belyayev 2014; Joly-Lopez and Bureau 2018) and WGD events (Hoffman et al. 2012) are potential triggers of morphological or adaptive key innovations and rapid diversification (Tank et al. 2015; Soltis and Soltis 2016). These aspects raise further questions regarding the origin and evolutionary mechanisms giving rise to the diversity of magnoliids (Sauquet and Magallón 2018). The slow but regular reduction in population size of *Annona muricata* is compatible with the Quaternary contraction of tropical regions in several parts of the world, and suggests that the soursop could be severely affected by climate changes, as may other tropical taxa. Contrary to the situation in most crop plants (Eyre-Walker et al. 1998), this reduction did not result from a genetic bottleneck during domestication. However, the very low heterozygosity in soursops could make future genetic improvement difficult, and will likely require outcrossing with wild relatives (Zamir 2001).

The soursop genome is smaller (657Mb) than *Liriodendron chinense* (1.75Gb) or *Cinnamomum kanehirae* (824Mb), and it displays the same chromosomes number (7) as that reconstructed for the ancestor of angiosperms (Badouin et al. 2017). Crucially, both *L. chinense* (19 chromosomes) and *C. kanehirae* (12 chromosomes) show clear signs (Suppl. Fig. 2) of lineage specific whole genome duplication (WGD) and chromosomal rearrangement events occurring after their branching from their shared MRCA with monocots or eudicots.

The comparative genomic analyses using three magnoliid genomes (*Annona, Cinnamomum* and *Liriodendron*) confirm some of the findings presented in these other studies, but also raise important analytical considerations in such analyses.

Especially, our results suggest that evidence for WGD in other magnoliids represent events that occurred subsequent, not prior, to their divergence, demonstrating the importance of increased representation in phylogenomic analyses of older lineages to improve our understanding of the early diversification of angiosperms. The soursop genome further highlights the limitations of using one species as a representative for a group as diverse as the magnoliids. Using one species per lineage makes it difficult to distinguish specific and shared WGD, especially in case of ancient events (Tiley et al. 2018). Ks distributions are useful to characterize specific WGD events, but for numbers of WGD events should be interpreted with caution (Tiley et al. 2018). Despite using a more robust method which avoids overfitting of a component-rich lognormal model to our distribution, we did not find clear evidence in favor of a given number of components (i.e. WGD events) in *Cinnamomum*.

A further emerging concern for phylogenomic analyses is the apparent unfavorable reciprocity between the number of taxa involved and the number of retrieved orthologs. We show that the number of orthologs used to perform phylogenomic reconstruction strongly impacts the retrieved topology, with too few genes also potentially resulting in the reconstruction of erroneous relationships.With the development of new methods for both sequencing (e.g. Nanopore, Hi-C) and analyses [e.g. paleo-karyotypes (Murat et al. 2014)], it becomes feasible to get high quality, chromosome-scale assemblies that can address more complex evolutionary questions. Our current study is limited by the unknown arrangement of scaffolds relative to each other, hampering our ability to reconstruct a high-resolution genomic comparison of the landscape of soursop with other magnoliids. Early angiosperms divergences also cannot be fully resolved without taking into account the other basal angiosperms lineages, including the elusive Chloranthales.

Here, we present results based on three magnoliids genomes, a very small part of the clade’s diversity. To improve our understanding of relationships within the group and structural rearrangements at and below the level of the genome, it will be vital to add representatives from other divergent lineages (e.g. Piperales, Cannelales), as well as other lineages in Magnoliales and Laurales.

The soursop genome, in addition to *Liriodendron* and *Cinnamomum*, will be an exceptional resource not only for the scientific community but also for breeders (avocado, *Annona* species, pepper, *Magnolia*, etc). Indeed, genomes give positional information that transcriptomes are unable to provide, allowing more sensitive and robust delineation of the WGD from tandem duplications (e.g. through synteny graphs) (Tiley et al. 2018), as well as allowing breeders to use linkage disequilibrium estimation in their programs (Barabaschi et al. 2015).

## Materials & Methods

### Plant materials

We identified a mature *Annona muricata* tree in Xishuangbanna Botanical Garden, located in the southern part of Yunnan province, China (21°55′41.8″N 101°15′31.7″E). The sampled tree was vouchered and tissues immediately frozen in liquid nitrogen upon collecting until sequencing experiments were performed.

### Genomic DNA extraction and library preparation

The libraries were subjected to the paired-end 150bp sequencing on the Illumina HiSeq 2500 platform. Approximately 900 millions reads were generated from Illumina libraries with different inserts sizes (250bp, 350bp) to provide a first estimation of the genome size, GC content, heterozygosity rate and repeat content. Genomic DNA was isolated from the leaves of *A. muricata*. For a 20-kb insert size library, at least 20 μg of sheared DNA was required. SMRTbell template preparation involved DNA concentration, damage repair, end repair, ligation of hairpin adapters, and template purification, and used AMPure PB Magnetic Beads. Finally, the sequencing primer was annealed and sequencing polymerase was bound to SMRTbell template. The instructions specified as calculated by the RS Remote software were followed. We carried out 20-kb single-molecule real-time DNA sequencing by PacBio and sequenced the DNA library on the PacBio RS II platform, yielding about 37Gb PacBio data (read quality ≥ 0.80, mean read length ≥ 7 Kb)

### 10X Genomics and Bionano library preparation and sequencing

DNA sample preparation, indexing, and barcoding were done using the GemCode Instrument from 10X Genomics. About 0.7 ng input DNA with 50 kb length was used for GEM reaction procedure during PCR, and 16-bp barcodes were introduced into droplets. Then, the droplets were fractured following the purifying of the intermediate DNA library. Next, we sheared DNA into 500 bp for constructing libraries, which were finally sequenced on the Illumina HiSeq X.

This sequencing strategy provided sequencing depths of 163X, 46X, 225X and 120X for Illumina, PacBio, 10X Genomics and Bionano libraries sequencing, respectively.

### RNA extraction and library preparation

Total RNA was extracted using the RNAprep Pure Plant Kit and genomic DNA contamination was removed using RNase-Free DNase I (both from Tiangen). The integrity of RNA was evaluated on a 1.0% agarose gel stained with ethidium bromide (EB), and its quality and quantity were assessed using a NanoPhotometer spectrophotometer (IMPLEN) and an Agilent 2100 Bioanalyzer (Agilent Technologies). As the RNA integrity number (RIN) was greater than 7.0 for all samples, they were used in cDNA library construction and Illumina sequencing, which was completed by Beijing Novogene Bioinformatics Technology Co., Ltd. The cDNA library was constructed using the NEBNext Ultra RNA Library Prep Kit for Illumina (NEB) and 3 μg RNA per sample, following the manufacturer’s recommendations. The PCR products obtained were purified (AMPure XP system) and library quality was assessed on the Agilent Bioanalyzer 2100 system. Library preparations were sequenced on an Illumina HiSeq 2000 platform, generating 100-bp paired-end reads.

### Estimation of genome size and heterozygosity

The k-mer frequency analysis, also known as k-mer spectra, is an efficient assembly-independent method for accurate estimation of genomic characteristics (genome size, repeat content, heterozygous rate). The k-mers refer to all the possible subsequences (of length k) from a read. We estimated the genome size based on the 17-mer frequency of Illumina short reads using the formula: genome size = (total number of 17-mer)/ (position of peak depth). We obtained an estimate of 799.11 Mb.

### Genome assembly

We used ALLPATHS-LG (Gnerre et al. 2011) and obtained a preliminary assembly of *A. muricata* with a scaffold N50 size of 19,908 kb and corresponding contig N50 size of 8.26 Kb. We used PBjelly (English et al. 2012) to fill gaps with PacBio data. The options were “<blasr>-minMatch 8 -sdpTupleSize 8 -minPctIdentity 75 -bestn 1 -nCandidates 10 -maxScore -500 -nproc 10 -noSplitSubreads</blasr>” for the protocol.xml file. Then, we used Pilon (Walker et al. 2014) with default settings to correct assembled errors. For the input BAM file, we used BWA to align all the Illumina short reads to the assembly and SAMtools to sort and index the BAM file.

We used BWA mem to align the 10X Genomics data to the filled gaps assembly using default settings. Then, we used fragScaff (https://sourceforge.net/projects/fragscaff/) for scaffolding.

### Transcriptome assembly

RNA sequencing (RNA-Seq) libraries were constructed using the NEBNext mRNA Library Prep Master Mix Set for Illumina following the manufacturer’s instructions. The libraries were subjected to paired-end 150 bp sequencing on the Illumina HiSeq 2000 platform. Raw reads were first filtered by removing those containing undetermined bases (‘N’) or excessive numbers of low-quality positions (>10 positions with quality scores <10). Then the high-quality reads were mapped to the U+0078^2^*A. muricata* genome using Tophat (v2.0.9)66 with the parameters of ‘-p 10 -N 3 --read-edit-dist 3 -m 1 -r 0 --coverage-search --microexon-search’.

### Genome annotation

Transposable elements in the genome assembly were identified both at the DNA and protein level. We used RepeatModeler to develop *de novo* transposable element library. RepeatMasker (Smit et al. 2017) was applied for DNA-level identification using Repbase and *de novo* transposable element library. At protein level, RepeatProteinMask was used to conduct WU-BLASTX searches against the transposable element protein database. Overlapping transposable elements belonging to the same type of repeats were integrated together.

Protein coding genes were predicted through combination of homology-based prediction, *de novo* predictions and transcriptome based predictions:

- Structural annotation of protein coding genes and protein domains was performed by aligning the protein sequences to the soursop genome by using TblastN with an E-value cutoff by 1E-5. The blast hits were conjoined by solar. For each blast hit, Genewise was used to predict the exact gene structure in the corresponding genomic regions.
- Five *ab initio* gene prediction programs, including Augustus (http://bioinf.uni-greifswald.de/augustus/), Genscan (http://genes.mit.edu/GENSCAN.html), GlimmerHMM (ftp://ccb.jhu.edu/pub/software/glimmerhmm), Geneid (http://genome.crg.es/geneid.html) and SNAP (https://github.com/KorfLab/SNAP), were used to predict coding genes on the repeat-masked genomes.
- Finally, RNA-seq data were mapped to genome using Tophat (Kim et al. 2013), and then cufflinks (http://cufflinks.cbcb.umd.edu/) was used to assemble transcripts to gene models.

All gene models predicted from the above three approaches were combined by EVidenceModeler (EVM) (Haas et al. 2008) into a non-redundant set of gene structures. Then we filtered out low quality gene models: (1) coding region lengths of ≤150 bp, (2) supported only by ab initial methods and with FPKM<1.

Functional annotation of protein coding genes was evaluated by BLASTP (evalue1E-05) instead of two integrated protein sequence databases – SwissProt and TrEMBL. The annotation information of the best BLAST hit, which is derived from database, was transferred to our gene set. Protein domains were annotated by searching InterPro (https://www.ebi.ac.uk/interpro/) and Pfam (https://pfam.xfam.org/) databases, using *InterProScan* and *Hmmer*, respectively. Gene Ontology (GO) terms for each gene were obtained from the corresponding InterPro or Pfam entry. The pathways, in which the gene might be involved, were assigned by blast against the KEGG database (https://www.genome.jp/kegg/), with an E-value cutoff of 1E-05.

The tRNA genes were identified by tRNAscan-SE (Lowe and Eddy 1996) with eukaryote parameters. The rRNA fragments were predicted by aligning to *Arabidopsis thaliana* and *Oryza sativa* template rRNA sequences using BlastN at E-value of 1E-10. The miRNA and snRNA genes were predicted using INFERNAL by searching against the Rfam database (http://rfam.xfam.org/).

### Population size changes inference

We used PSMC (Liu and Hansen 2017) to infer the variation in population size of the soursop based on the observed heterozygosity in the diploid genome. As PSMC was shown to performed reliably for scaffolds >100kb, we removed shorter scaffolds from the assembly. 312 scaffolds > 100kb were kept, totalizing 646.64 Mb (98.46 percent of the total assembly). We assume a generation time of 15 years (Collevatti et al. 2014) and a per-generation mutation rate of 7×10-9. PSMC was otherwise conducted using default parameters.

### Coalescence phylogenetic analyses

We selected the following 12 representative plant species (Supplementary Table XX): *C. canephora, H. annuus, A. hypochondriacus* (in Super-Asterids), *V. vinifera, A. thaliana, Q. robur* (in Rosids), *M.acuminata, O. sativa* (Monocots), *N. nucifera* and *A. coerulea*, plus two recently published magnoliids genomes (*Cinnamomum kanehirae* and *Liriodendron chinense*). We reconstructed phylogenetic trees using the same method with three datasets:

- orthologs identified in the 12 representative plant species, maximizing the number of species;
- orthologs identified in the quartet (*Annona muricata, Arabidopsis thaliana, Amborella trichopoda* and *Oryza sativa*), maximising the number of loci;
- orthologs identified in the previous quartet, plus *Cinnamomum kanehirae* and *Liriodendron chinense*, maximizing the magnoliids representation;

For each dataset, we used OrthoMCL (Li et al. 2003) to define gene family clusters among different species. An all-against-all BLASTP was first applied to determine the similarities between genes in all genomes at the E-value threshold of 1e-7. Then the Markov clustering (MCL) algorithm implemented in OrthoMCL was used to group orthologs and paralogs from all input species with an inflation value (-I) of 1.5. The sequences from each family containing one single ortholog for all species were aligned using mafft and a phylogenetic reconstructed using PhyML (Guindon et al. 2010) with a GTR+I+G model with 4 substitution categories and 100 bootstrap replicates. The species trees were then built for each dataset with MP-est, NJ-st and STAR as implemented in the Species Tree Webserver (Shaw et al. 2013) and ASTRAL-III (Zhang et al. 2018) using default parameters.

### Gene family expansion analysis

The species tree from the dataset maximizing the magnoliids representation was used to analyze the Gene family expansion/contraction along the branch of the species tree with CAFE (De Bie et al. 2006), with correction for potential assembly and annotations mistakes. We dated the ASTRAL-III tree with treePL (Smith and O’Meara 2012) and several nodes calibrations (root max age = 200Ma (Massoni et al. 2015); divergence Monocots/Eudicots min-max age: 160-195Ma (Foster et al. 2016); divergence Magnoliales/Laurales min-max age: 121-162 Ma (Massoni et al. 2015); Magnoliales crown min-max age: 114-146Ma (Massoni et al. 2015)). One representative sequence from each family was used for downstream annotation as above. GO-terms were retrieved for annotated in expanded families using PANTHER (http://www.pantherdb.org/).

### Identification of WGD events in *A. muricata* and during early angiosperm divergence

*K*S-based age distributions were constructed using wgd (Zwaenepoel and Van de Peer 2019). In brief, the paranome of magnoliids and one-vs-one orthologs comparisons were constructed by performing all-against-all protein sequence similarity searches using BLASTP with an *E* value cutoff of 1 × 10−10, after which gene families were built with mcl (Altschul et al. 1997; Stijn Marinus van Dongen 2000). Each gene family was aligned using mafft (Katoh and Standley 2013) and *K*S estimates for all pairwise comparisons within a gene family were obtained through maximum likelihood estimation using CODEML (Kohlhase 2006) of the PAML package (Yang 2007). Gene families were then subdivided into subfamilies for which KS estimates between members did not exceed a value of 5. To correct for the redundancy of *K*S values (a gene family of *n* members produces *n*(*n*−1)/2 pairwise *K*S estimates for *n*−1 retained duplication events), a phylogenetic tree was constructed for each subfamily using FastTree (Price et al. 2009) under default settings. For each duplication node in the resulting phylogenetic tree, all *m K*S estimates between the two child clades were added to the *K*S distribution with a weight of 1/*m* (where *m* is the number of *K*S estimates for a duplication event), so that the weights of all *K*S estimates for a single duplication event summed to one.

### Phylogenomic incongruence analyses

To investigate the patterns of incongruence among loci in reconstructing the early history of angiosperms, we analyzed individually each of the 689 orthologous genes found in 12 selected angiosperms. For each gene, orthologous sequences were aligned as described above, then PhyML was used with the WAG model and SH-like confidence estimation with *A. trichopoda* set as outgroup. The resulting trees were sorted using the sortTrees function in R to identify trees with a given topology at a specific node [i.e. Amur+Eu dicotyledons, Amur+Monocotyledons or Amur,(Eudicotyledons+Monocotyledons)], according to the support at this node. We then extracted the GO-terms annotations for the genes supporting each topology with each support using Panther (Mi et al. 2018) and custom scripts.

In order to increase the number of orthologous sequences used and to assess the influence of the data matrix configuration (“less genes x more species” vs “more gens x less species”), we generated orthologous sequences set for a restricted number of species representative of the major clades in angiosperms (namely Magnoliids – *A. muricata*, Monocots – *O. sativa*, Dicots – *A. thaliana*, and *A. trichopoda*). This set of genes contained 2426 orthologous coding genes found in the 4 species in single copies, and was analysis in the same way than describe above for the 689 loci dataset.

We then explore the genomic landscape of incongruence loci by comparing the scaffold containing genes supporting a given topology for the second dataset. Heatmaps for several parameters (i.e. the number of genes, their density and their relative abundance compared to the background (the entire set of orthologs) were generated using the heatmap.2 function from the gplots package in R (R Development Core Team 2010).

A linear regression (“lm” function in R) was used to confirm the portion of genes supporting a given topology was independent of the scaffold length (p-values > 0.2 for each comparison).

## Acknowledgements

Genome sequencing, assembly and annotation were conducted by the Novogene Bioinformatics Institute, Beijing, China; mutual contract No. NHT161060. This work was supported by funding through the Guangxi Province One Hundred Talent program and Guangxi University to JSS, and the China Postdoctoral Science Foundation (grant number 2015M582481 and 2016T90822) to DDH. The basis for this manuscript was laid down during the 2015 dialogue seminar on Annonaceae under the Joint Scientific Thematic Research Programme (JSTP) funded by the Netherlands Organisation for Scientific Research and the Chinese Academy of Sciences (grant number 045.011.020). TLPC was supported by the Agence Nationale de la Recherche (grant AFRODYN: ANR-15-CE02-0002-01) and acknowledges the IRD itrop South Green Platform at IRD montpellier for providing HPC resources that have contributed to the research results reported within this paper. MDP is supported by the Heisenberg programme of the Deutsche Forschungsgemeinschaft (PI 1169/3-1). The authors report no conflict of interests.

## Code availability

All custom scripts used are available from the corresponding author upon request.

## Data availability

Genome sequences and whole-genome assembly of *A. muricata* and whole transcriptomes have been submitted to the National Center for Biotechnology Information (NCBI) database under BioProject PRJEB30626 (pending). All other data are available from the corresponding authors upon reasonable request.

## Supplementary Figures captions

**Supplementary figure 1.** K-mer distribution analysis for genome size and heterozygosity estimation.

**Supplementary Figure 2.** Ks plots for different pairs of comparisons. Histograms show the frequency of the pairwise comparisons for the given value of Ks, solid curves are densities of the gaussian mixture models, each dashed curve corresponds to a separate Gaussian component. a. pairwise comparison of Annona muricata paralogs; b. pairwise comparison of Cinnamomum kanehirae paralogs; c. pairwise comparison of Liriodendron chinense paralogs; d: pairwise comparisons of orthologs between Annona and Cinnamomum; e: pairwise comparisons of orthologs between Annona and Liriodendron; f: pairwise comparisons of orthologs between Liriodendron and Cinnamomum; g: pairwise comparisons of orthologs between Annona and Amborella. Insets in a,b and c show the Bayesian Information Criterion (BIC, top) and Akaike Information Criterion (AIC, bottom) according to the number of components considered in the mixture, arrows indicate the number of components shown on the figure used for fitting the Ks histogram; red and blue arrows in b correspond to two and three components used to fit the Ks histogram, respectively. Black vertical line in d, e and f shows the mode of the Annona vs Liriodendron Ks values distribution.

**Supplementary Figure 3.** Gene prediction support. Number of predicted genes supported by RNA-seq transcripts (rna_0.5), homology to known proteins (homolog_0.5) or ab initio inference (denovo_0.5).

**Supplementary Figure 4.** Coalescent analyses of the 689 loci (a, b, c), 2426 loci (e, f, g, h) and 1578 loci (i, j, k, l) datasets using MP-est (a, e, i), NJ-st (b, f, j), STAR (c,g,k) and ASTRAL-III (d, h, l) algorithms. MP-est, NJ-st and STAR analyses were performed using the STRAW webserver 6. Branch lengths in a and e are in coalescent units. The branch leading to Annona muricata is shown in red. Bootstrap values indicated at nodes when available.

**Supplementary Figure 5.** PCA plot comparing the annotations for genes supporting the SET, SAT and SMT with different supports (SH-like values <0.50, 0.50<SH-like values <0.70, 0.70<SH-like values <0.95, SH-like values >0.95) based on GO-term annotations content. PC1 and PC2 are shown. AnnMonocot: SMT; Dicots: SET; basalAnnona: SAT; background: general annotations content of the soursop genome. a. Biplot of the datasets, with contribution of the variable to the axes shown as blue arrows. Points colors according to cos 2. b. Colors according to the supported topology. Ellipses indicate 95% confidence interval for belonging to a group based on annotations content.

**Supplementary Figure 6.** Genomic landscape of incongruent phylogenomic signal. Colors according to row Z-scores (a statistical estimation of the spread of the value for each gene compared to the mean). Each column represents a given topology; each line is a scaffold. a: number of gene supporting a given topology per scaffold; b: number of gene per base of scaffold; c: percentage of orthologs strongly supporting a given topology (relative to the total number of genes supporting this topology, i.e. the ratio “number of genes supporting the topology in the scaffold” / “total number of genes with support >0.70 for this topology in the dataset”) found in each scaffold; d: difference between the density of genes supporting a given topology and the density of orthologous genes in the scaffolds (aka the background).

**Supplementary Figure 7.** Plots of the number of genes supporting a given topology relative to the total number of genes in the scaffold (i.e. the background repartition of the 2426 orthologous loci) showing a higher proportion of genes supporting a basal Annona hypothesis (Main panel). Colors of linear regression and their slope p-values according according to the supported hypothesis green: genes supporting SAT; red: genes supporting SET; blue: genes supporting SMT. Left panel: density plot of the relative number of genes supporting a given topology. Bottom panel: density plot of the scaffold lengths. Inset: boxplot of the same data than the main panel without considering the scaffold length.

## Supplementary Tables captions

**Supplementary Table 1.** Overview of the sequences generated.

**Supplementary Table 2.** Assembly statistics.

**Supplementary Table 3.** To evaluate the quality of the genome assembly, we mapped reads from short insert size libraries back to the scaffolds by using BWA (http://bio-bwa.sourceforge.net/). The sequencing depth distribution follows a Poisson distribution, which indicates the uniformity of the genome sequencing process.

**Supplementary Table 4.** Classification of TEs content.

**Supplementary Table 5.** Characteristics of the annotated genes in 10 angiosperms species.

**Supplementary Table 6.** Overview of annotated genes per database.

**Supplementary Table 7.** CEGMA and BUSCO assessments of the genes annotations.

**Supplementary Table 8.** Number of trees supporting each topology for the 12 and 4 species dataset, respectively. Trees are classified according to the support (SH-like values) of the node supporting the topology.

**Supplementary Table 9.** Identification of functional biases among orthologous sequences in the 4 species dataset. Numbers of genes attributed to given GO-term among the orthologs supporting different topologies.

**Supplementary Table 10.** Statistics of the distribution of orthologs across scaffolds. Density is expressed in number of orthologs per base. The background is defined as the total dataset (2426 loci in 181 scaffolds). Differential density is expressed as the ratio of the density of genes supporting a given topology over the background density of orthologs.

## References

Altschul SF, Madden TL, Schäffer AA, Zhang J, Zhang Z, Miller W, Lipman DJ. 1997. Gapped BLAST and PSI-BLAST: A new generation of protein database search programs. Nucleic Acids Res. 25:3389–3402.

Badouin H, Gouzy J, Grassa CJ, Murat F, Staton SE, Cottret L, Lelandais-Brière C, Owens GL, Carrère S, Mayjonade B, et al. 2017. The sunflower genome provides insights into oil metabolism, flowering and Asterid evolution. Nature 546:148–152.

Barabaschi D, Tondelli A, Desiderio F, Volante A, Vaccino P, Valè G, Cattivelli L. 2015. Next generation breeding. Plant Sci. 242:3–13.

Bell CD, Soltis DE, Soltis PS. 2010. The age and diversification of the angiosperms re-revisited. Am. J. Bot. 97:1296–1303.

Belyayev A. 2014. Bursts of transposable elements as an evolutionary driving force. J. Evol. Biol. 27:2573–2584.

De Bie T, Cristianini N, Demuth JP, Hahn MW. 2006. CAFE: A computational tool for the study of gene family evolution. Bioinformatics 22:1269–1271.

Chatrou LW, Erkens RHJ, Richardson JE, Saunders RMK, Fay MF. 2012. The natural history of Annonaceae. Bot. J. Linn. Soc. 169:1–4.

Chaw SM, Liu YC, Wu YW, Wang HY, Lin CYI, Wu CS, Ke HM, Chang LY, Hsu CY, Yang HT, et al. 2019. Stout camphor tree genome fills gaps in understanding of flowering plant genome evolution. Nat. Plants 5:63.

Chen J, Hao Z, Guang X, Zhao C, Wang P, Xue L, Zhu Qihui, Yang Linfeng, Sheng Y, Zhou Y, et al. 2019. <>Liriodendron</i> genome sheds light on angiosperm phylogeny and species–pair differentiation. Nat. Plants 5:18.

Collevatti RG, Telles MPC, Lima JS, Gouveia FO, Soares TN. 2014. Contrasting spatial genetic structure in Annona crassiflora populations from fragmented and pristine savannas. Plant Syst. Evol. 300:1719–1727.

Doebley JF, Gaut BS, Smith BD. 2006. The Molecular Genetics of Crop Domestication. Cell 127:1309–1321

Edwards S V. 2009. Is a new and general theory of molecular systematics emerging? Evolution (N. Y). [Internet] 63:1–19. Available from: https://dash.harvard.edu/bitstream/handle/1/26514972/Edwards_2009_Commentary_accepted_version.pdf?sequence=1

Edwards S V., Xi Z, Janke A, Faircloth BC, McCormack JE, Glenn TC, Zhong B, Wu S, Lemmon EM, Lemmon AR, et al. 2016. Implementing and testing the multispecies coalescent model: A valuable paradigm for phylogenomics. Mol. Phylogenet. Evol. [Internet] 94:447–462. Available from: https://www.sciencedirect.com/science/article/pii/S1055790315003309

English AC, Richards S, Han Y, Wang M, Vee V, Qu J, Qin X, Muzny DM, Reid JG, Worley KC, et al. 2012. Mind the Gap: Upgrading Genomes with Pacific Biosciences RS Long-Read Sequencing Technology. PLoS One [Internet] 7:e47768. Available from: https://doi.org/10.1371/journal.pone.0047768

Eyre-Walker A, Gaut RL, Hilton H, Feldman DL, Gaut BS. 1998. Investigation of the bottleneck leading to the domestication of maize. Proc. Natl. Acad. Sci. U. S. A. 95:4441–4446.

Foster CSP, Sauquet H, Van Der Merwe M, McPherson H, Rossetto M, Ho SYW. 2016. Evaluating the impact of genomic data and priors on Bayesian estimates of the angiosperm evolutionary timescale. Syst. Biol. 66:338–351.

Freitas L, Mello B, Schrago CG. 2018. Multispecies coalescent analysis confirms standing phylogenetic instability in Hexapoda. J. Evol. Biol. [Internet] 31:1623–1631. Available from: http://doi.wiley.com/10.1111/jeb.13355

Gentry AH. 1993. Four neotropical rainforests. New Haven: Yale University Press

Gnerre S, MacCallum I, Przybylski D, Ribeiro FJ, Burton JN, Walker BJ, Sharpe T, Hall G, Shea TP, Sykes S, et al. 2011. High-quality draft assemblies of mammalian genomes from massively parallel sequence data. Proc. Natl. Acad. Sci. 108:1513–1518.

Guindon S, Dufayard JF, Lefort V, Anisimova M, Hordijk W, Gascuel O. 2010. New algorithms and methods to estimate maximum-likelihood phylogenies: Assessing the performance of PhyML 3.0. Syst. Biol. 59:307–321.

Haas BJ, Salzberg SL, Zhu W, Pertea M, Allen JE, Orvis J, White O, Robin CR, Wortman JR. 2008. Automated eukaryotic gene structure annotation using EVidenceModeler and the Program to Assemble Spliced Alignments. Genome Biol. 9:1.

Hoffman FG, Opazo JC, Storz JF. 2012. Whole-genome duplications spurred the functional diversification of the globin gene superfamily in vertebrates. Mol. Biol. Evol. 29:303–312.

Initiative A genome. 2000. Analysis of the genome sequence of the flowering plant Arabidopsis thaliana. Nature 408:796.

Jarvis ED, Mirarab S, Aberer AJ, Li B, Houde P, Li C, Ho SYW, Faircloth BC, Nabholz B, Howard JT, et al. 2014. Whole-genome analyses resolve early branches in the tree of life of modern birds. Science (80-.). 346:1320–1331.

Joly-Lopez Z, Bureau TE. 2018. Exaptation of transposable element coding sequences. Curr. Opin. Genet. Dev. [Internet] 49:34–42. Available from: https://www.sciencedirect.com/science/article/pii/S0959437X17301582

Katoh K, Standley DM. 2013. MAFFT multiple sequence alignment software version 7: improvements in performance and usability. Mol. Biol. Evol. [Internet] 30:772–780. Available from: http://www.pubmedcentral.nih.gov/articlerender.fcgi?artid=3603318&tool=pmcentrez&rendertype=abstract

Kim D, Pertea G, Trapnell C, Pimentel H, Kelley R, Salzberg SL. 2013. TopHat2: accurate alignment of transcriptomes in the presence of insertions, deletions and gene fusions. Genome Biol. 14:R36.

Kohlhase M. 2006. CodeML: an open markup format the content and presentatation of program code. Available from: https://svn.omdoc.org/repos/codeml/doc/spec/codeml

Li L, Stoeckert CJ, Roos DS. 2003. OrthoMCL: Identification of ortholog groups for eukaryotic genomes. Genome Res. 13:2178–2189.

Liu S, Hansen MM. 2017. PSMC (pairwise sequentially Markovian coalescent) analysis of RAD (restriction site associated DNA) sequencing data. Mol. Ecol. Resour. 17:631–641.

Lowe TM, Eddy SR. 1996. TRNAscan-SE: A program for improved detection of transfer RNA genes in genomic sequence. Nucleic Acids Res. 25:955–964.

Magallón S, Gómez-Acevedo S, Sánchez-Reyes LL, Hernández-Hernández T. 2015. A metacalibrated time-tree documents the early rise of flowering plant phylogenetic diversity. New Phytol. [Internet] 207:437–453. Available from: http://doi.wiley.com/10.1111/nph.13264

Massoni J, Couvreur TLP, Sauquet H. 2015. Five major shifts of diversification through the long evolutionary history of Magnoliidae (angiosperms) Phylogenetics and phylogeography. BMC Evol. Biol. 15:49.

Massoni J, Forest F, Sauquet H. 2014. Increased sampling of both genes and taxa improves resolution of phylogenetic relationships within Magnoliidae, a large and early-diverging clade of angiosperms. Mol. Phylogenet. Evol. 70:84–93.

Mi H, Muruganujan A, Ebert D, Huang X, Thomas PD. 2018. PANTHER version 14: more genomes, a new PANTHER GO-slim and improvements in enrichment analysis tools. Nucleic Acids Res. 47:D419–D426.

Moore MJ, Hassan N, Gitzendanner MA, Bruenn RA, Croley M, Vandeventer A, Horn JW, Dhingra A, Brockington SF, Latvis M, et al. 2011. Phylogenetic Analysis of the Plastid Inverted Repeat for 244 Species: Insights into Deeper-Level Angiosperm Relationships from a Long, Slowly Evolving Sequence Region. Int. J. Plant Sci. 172:541–558.

Moore MJ, Soltis DE, Burleigh JG, Bell CD, Soltis PS, Bell CD, Burleigh JG, Soltis DE. 2010. Phylogenetic analysis of 83 plastid genes further resolves the early diversification of eudicots. Proc. Natl. Acad. Sci. [Internet] 107:4623–4628. Available from: http://www.ncbi.nlm.nih.gov/entrez/query.fcgi?cmd=Retrieve&db=PubMed&dopt=Citation&list_uids=20176954

Murat F, Armero A, Pont C, Klopp C, Salse J. 2017. Reconstructing the genome of the most recent common ancestor of flowering plants. Nat. Genet. 49:490.

Murat F, Pont C, Salse J. 2014. Paleogenomics in Triticeae for translational research. Curr. Plant Biol. 1:34–39.

Nickrent DL, Soltis DE. 2006. A Comparison of Angiosperm Phylogenies from Nuclear 18S rDNA and rbcL Sequences. Ann. Missouri Bot. Gard. 82:208.

Oliver KR, McComb JA, Greene WK. 2013. Transposable elements: Powerful contributors to angiosperm evolution and diversity. Genome Biol. Evol. [Internet] 5:1886–1901. Available from: https://academic.oup.com/gbe/article-lookup/doi/10.1093/gbe/evt141

Price MN, Dehal PS, Arkin AP. 2009. Fasttree: Computing large minimum evolution trees with profiles instead of a distance matrix. Mol. Biol. Evol. 26:1641–1650.

Punyasena SW, Eshel G, McElwain JC. 2008. The influence of climate on the spatial patterning of Neotropical plant families. J. Biogeogr. 35:117.

Qiu YL, Li L, Wang B, Xue JY, Hendry TA, Li RQ, Brown JW, Liu Y, Hudson GT, Chen ZD. 2010. Angiosperm phylogeny inferred from sequences of four mitochondrial genes. J. Syst. Evol. 43:391–425.

R Development Core Team. 2010. R: A language and environment for statistical computing. Available from: http://www.r-project.org

Rainer H, Chatrou LW. 2014. AnnonBase: World species list of Annonaceae.

Richardson AO, Rice DW, Young GJ, Alverson AJ, Palmer JD. 2013. The “fossilized” mitochondrial genome of Liriodendron tulipifera: Ancestral gene content and order, ancestral editing sites, and extraordinarily low mutation rate. BMC Biol. 11:29.

Ruhfel BR, Gitzendanner MA, Soltis PS, Soltis DE, Burleigh JG. 2014. From algae to angiosperms-inferring the phylogeny of green plants (Viridiplantae) from 360 plastid genomes. BMC Evol. Biol. [Internet] 14:23. Available from: http://bmcevolbiol.biomedcentral.com/articles/10.1186/1471-2148-14-23

Sauquet H, von Balthazar M, Magallón S, Doyle JA, Endress PK, Bailes EJ, Barroso de Morais E, Bull-Hereñu K, Carrive L, Chartier M, et al. 2017. The ancestral flower of angiosperms and its early diversification. Nat. Commun. [Internet] 8:16047. Available from: http://www.nature.com/doifinder/10.1038/ncomms16047

Sauquet H, Magallón S. 2018. Key questions and challenges in angiosperm macroevolution. New Phytol. 219:1170–1187.

Shaw TI, Ruan Z, Glenn TC, Liu L. 2013. STRAW: Species TRee Analysis Web server. Nucleic Acids Res. 41:W238–W241.

Smit A, Hubley R, Green P. 2017. RepeatMasker Open-4.0.6 2013–2015. http://www.repeatmasker.org.

Smith SA, O’Meara BC. 2012. TreePL: Divergence time estimation using penalized likelihood for large phylogenies. Bioinformatics 28:2689–2690.

Soltis DE, Smith SA, Cellinese N, Wurdack KJ, Tank DC, Brockington SF, Refulio-Rodriguez NF, Walker JB, Moore MJ, Carlsward BS, et al. 2011. Angiosperm phylogeny: 17 genes, 640 taxa. Am. J. Bot. 98:704–730.

Soltis DE, Soltis PS. 2019. Nuclear genomes of two magnoliids. Nat. Plants 5:6.

Soltis PS, Soltis DE. 2016. Ancient WGD events as drivers of key innovations in angiosperms. Curr. Opin. Plant Biol. 30:159–165.

Song S, Liu L, Edwards S V., Wu S. 2012. Resolving conflict in eutherian mammal phylogeny using phylogenomics and the multispecies coalescent model. Proc. Natl. Acad. Sci. [Internet] 109:14942–14947. Available from: http://www.pnas.org/cgi/doi/10.1073/pnas.1211733109

Sonké B, Couvreur T. 2014. Tree diversity of the Dja Faunal Reserve, southeastern Cameroon. Biodivers. Data J. 2.

Stijn Marinus van Dongen. 2000. Graph clustering by flow simulation. Dr. Diss.

Tank DC, Eastman JM, Pennell MW, Soltis PS, Soltis DE, Hinchliff CE, Brown JW, Sessa EB, Harmon LJ. 2015. Nested radiations and the pulse of angiosperm diversification: Increased diversification rates often follow whole genome duplications. New Phytol. 207:454–467.

Tchouto MGP, Yemefack M, De Boer WF, De Wilde JJFE, Van Der Maesen LJG, Cleef AM. 2006. Biodiversity hotspots and conservation priorities in the Campo-Ma’an rain forests, Cameroon. Biodivers. Conserv. 15:1219–1252.

Tiley GP, Barker MS, Burleigh JG. 2018. Assessing the Performance of Ks Plots for Detecting Ancient Whole Genome Duplications. Genome Biol. Evol. 10:2882–2898.

Walker BJ, Abeel T, Shea T, Priest M, Abouelliel A, Sakthikumar S, Cuomo CA, Zeng Q, Wortman J, Young SK, et al. 2014. Pilon: An integrated tool for comprehensive microbial variant detection and genome assembly improvement. PLoS One [Internet] 9:e112963. Available from: https://doi.org/10.1371/journal.pone.0112963

Wickett NJ, Mirarab S, Nguyen N, Warnow T, Carpenter E, Matasci N, Ayyampalayam S, Barker MS, Burleigh JG, Gitzendanner MA, et al. 2014. Phylotranscriptomic analysis of the origin and early diversification of land plants. Proc. Natl. Acad. Sci. 111:E4859–E4868.

Yang Z. 2007. PAML 4: Phylogenetic analysis by maximum likelihood. Mol. Biol. Evol. 24:1586–1591.

Zamir D. 2001. Improving plant breeding with exotic genetic libraries. Nat. Rev. Genet. 2:983.

Zeng L, Zhang Q, Sun R, Kong H, Zhang N, Ma H. 2014. Resolution of deep angiosperm phylogeny using conserved nuclear genes and estimates of early divergence times. Nat. Commun. 5:4956.

Zhang C, Rabiee M, Sayyari E, Mirarab S. 2018. ASTRAL-III: Polynomial time species tree reconstruction from partially resolved gene trees. BMC Bioinformatics 19:153.

Zhang N, Zeng L, Shan H, Ma H. 2012. Highly conserved low-copy nuclear genes as effective markers for phylogenetic analyses in angiosperms. New Phytol. 195:923–937.

Zhu Q, Zheng X, Luo J, Gaut BS, Ge S. 2007. Multilocus analysis of nucleotide variation of Oryza sativa and its wild relatives: Severe bottleneck during domestication of rice. Mol. Biol. Evol. 24:875–888.

Zwaenepoel A, Van de Peer Y. 2019. wgd—simple command line tools for the analysis of ancient whole-genome duplications. Bioinformatics.

