## Supplementary figures and images for "The soursop genome and comparative genomics of basal angiosperms provide new insights on evolutionary incongruence"

### Supplementary Fig. 1

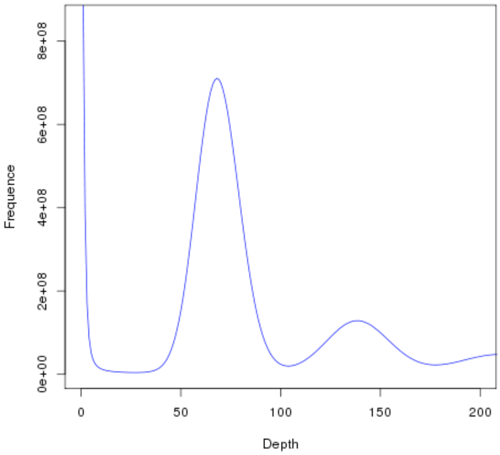

### Supplementary Fig. 2

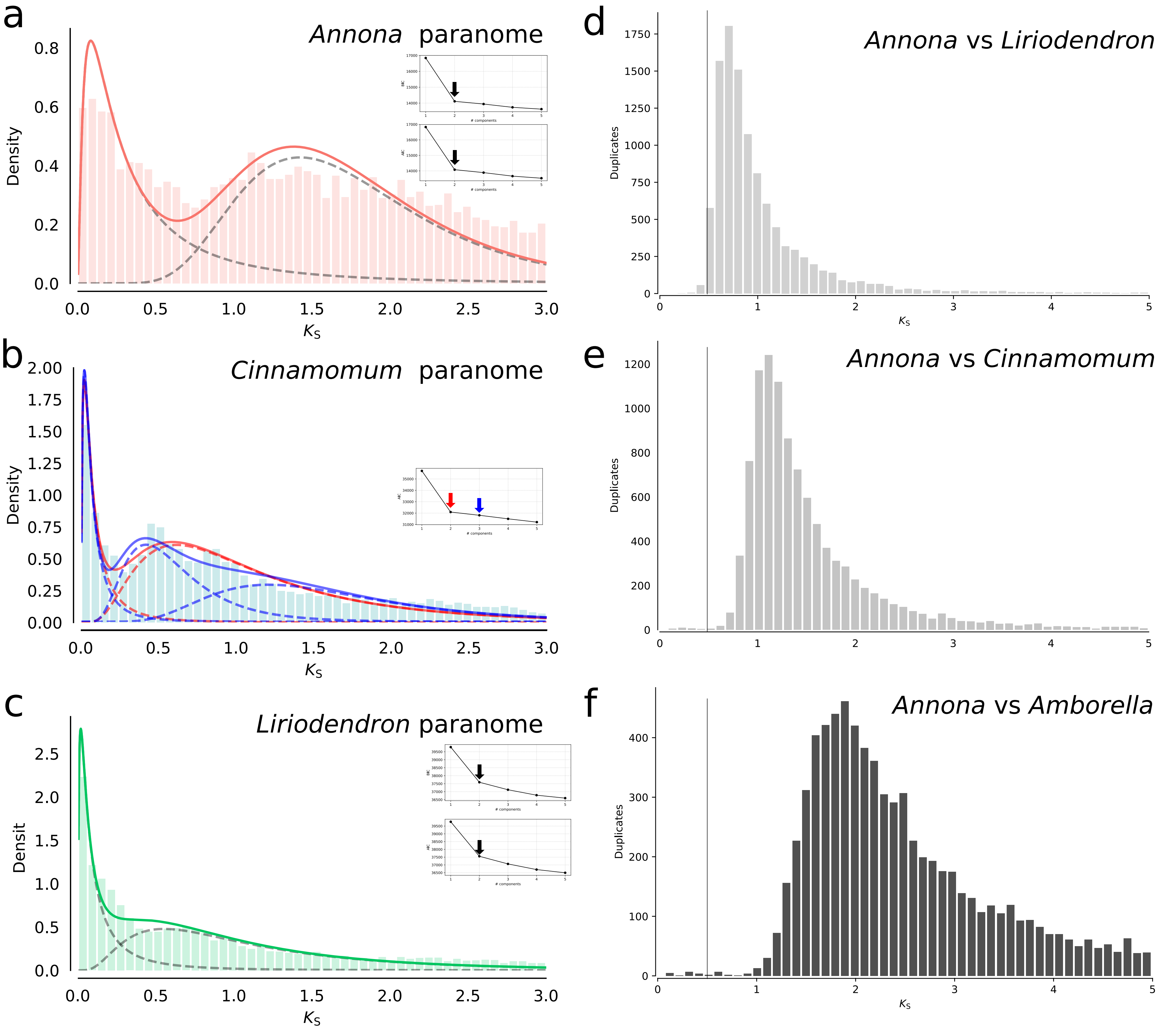

### Supplementary Fig. 3

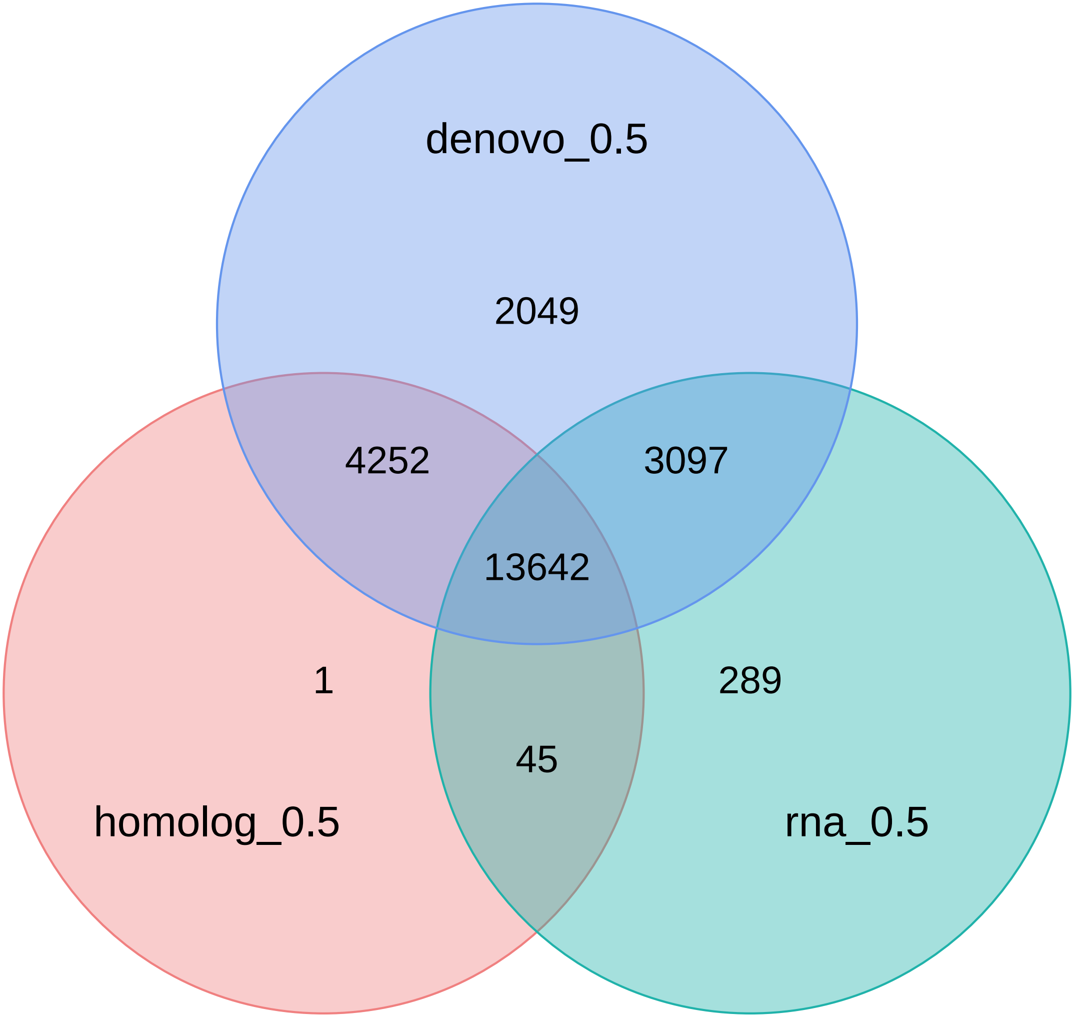


Evidence Support

### Supplementary Fig. 4

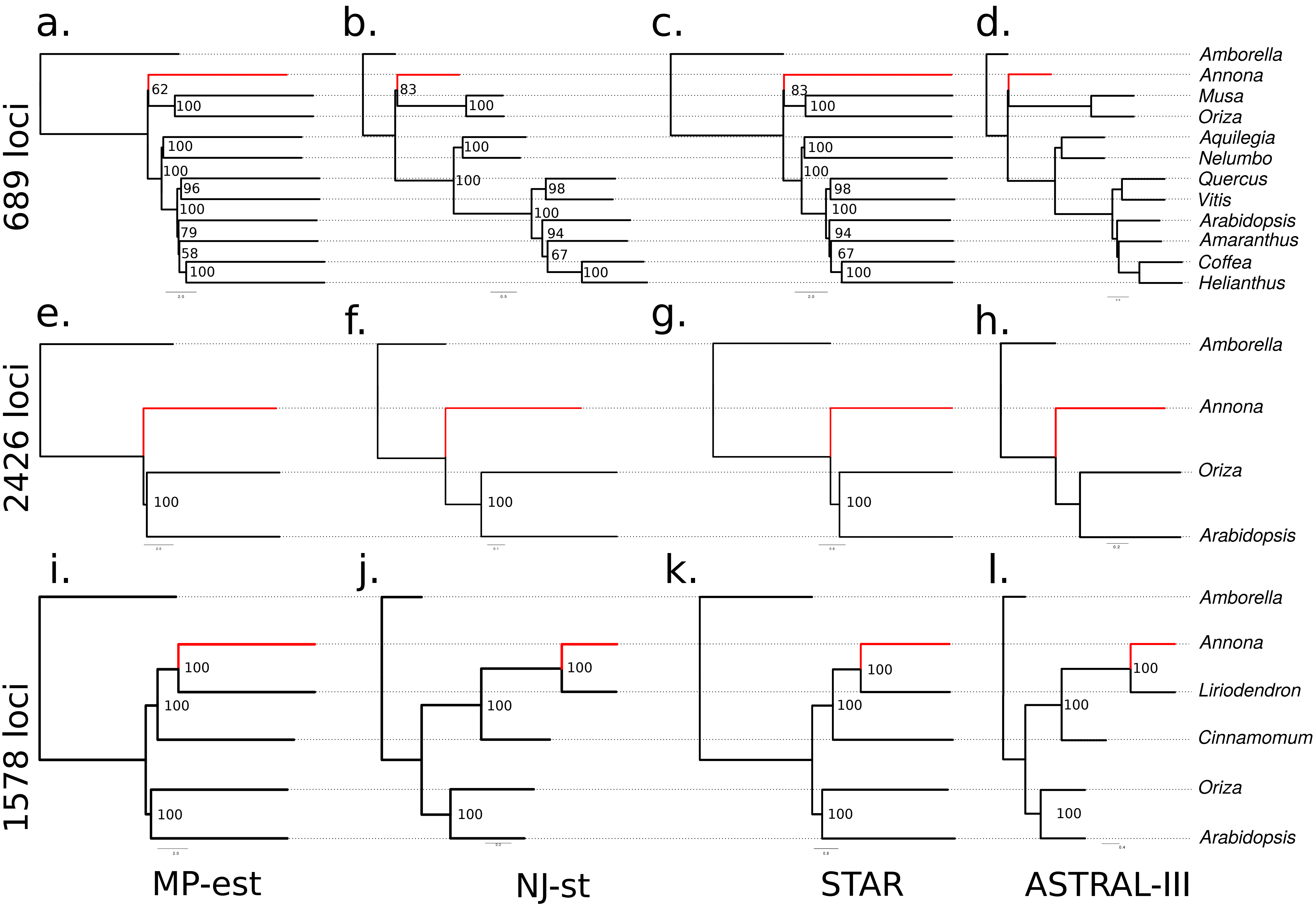

### Supplementary Fig. 5

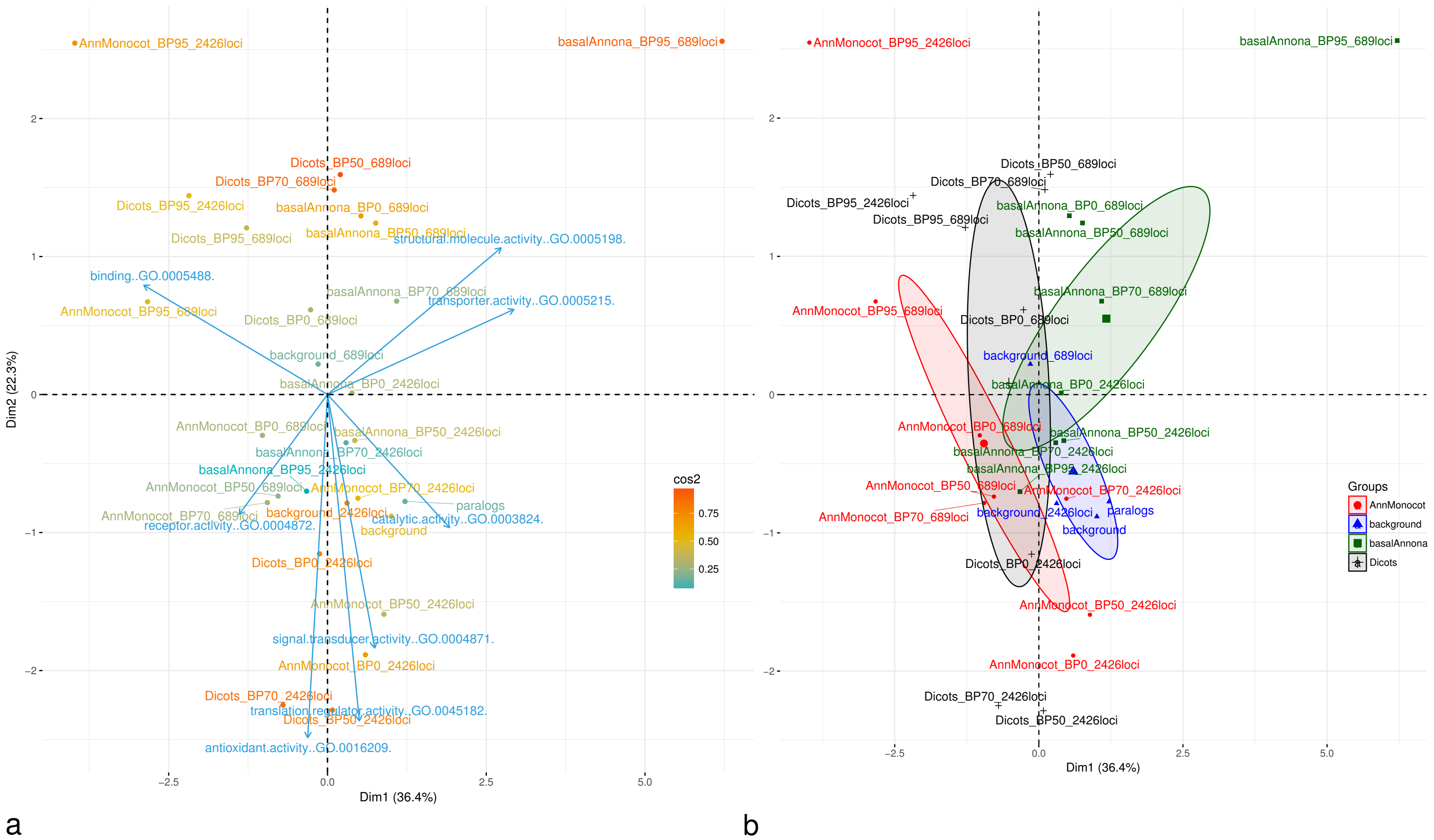

### Supplementary Fig. 6

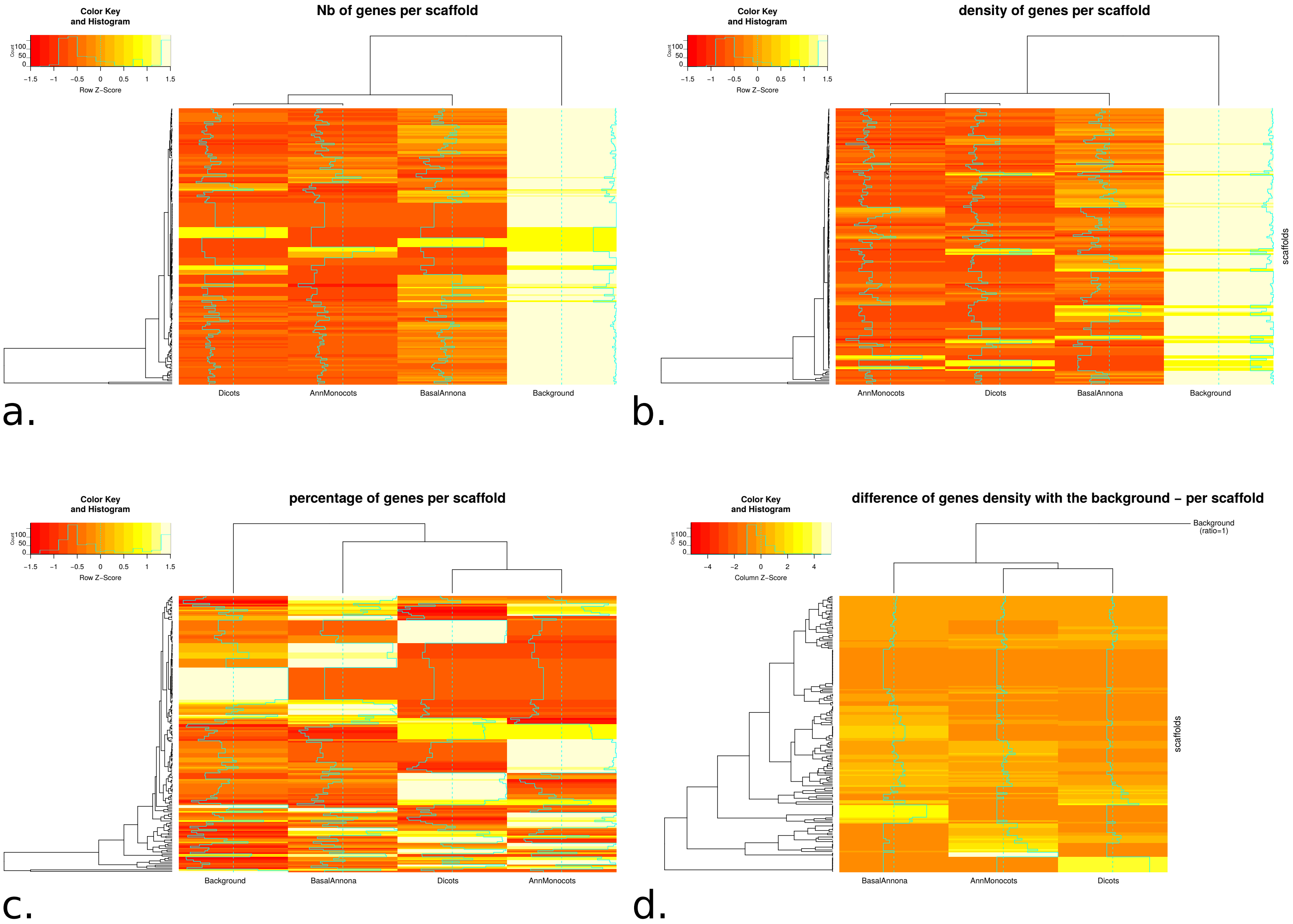

### Supplementary Fig. 7

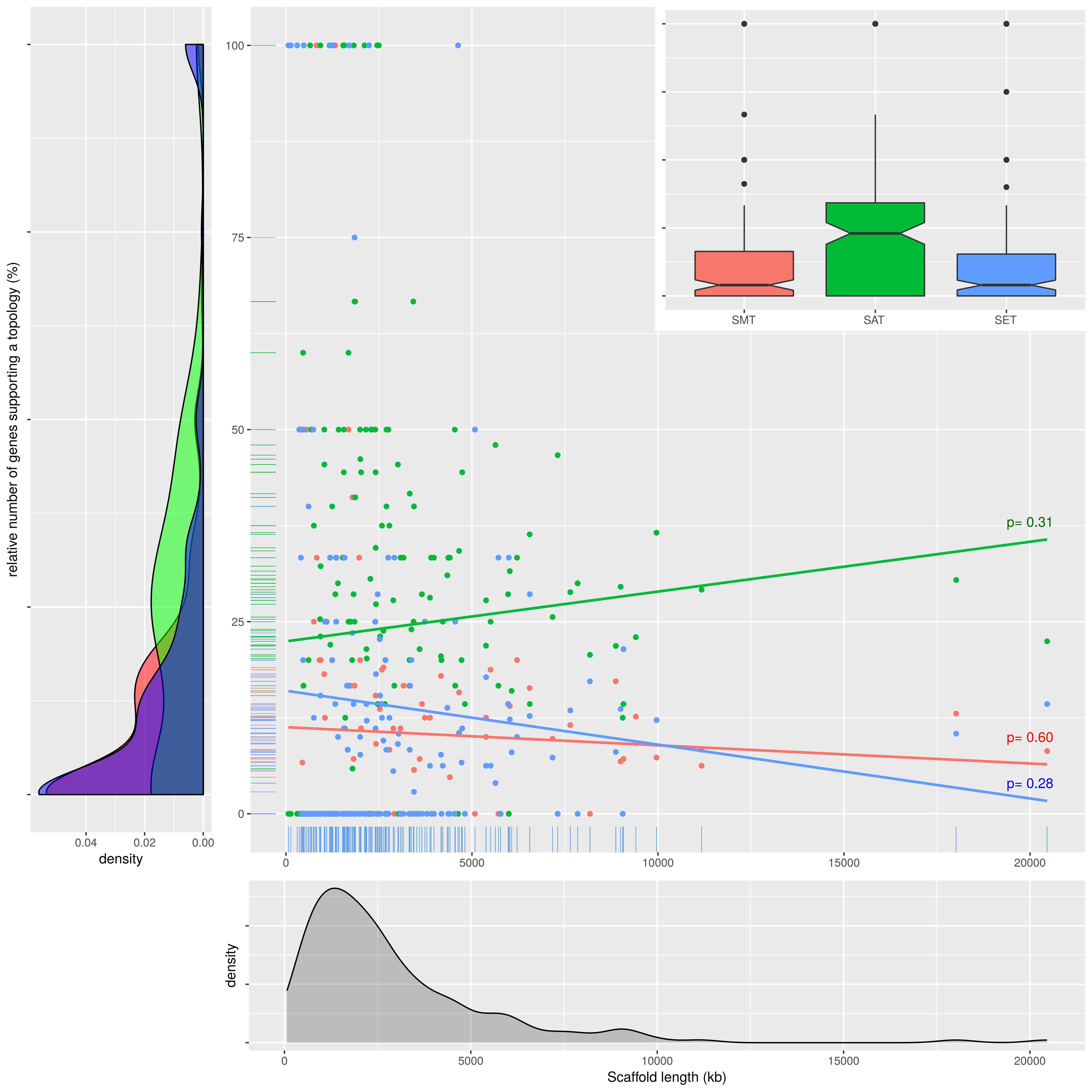
